# Integrated tear proteomics define the molecular blueprint of corneal epithelial repair

**DOI:** 10.1101/2025.07.17.665304

**Authors:** Nadege Feret, Marilou Decoudu, Jerome Vialaret, Christophe Hirtz, Vincent Daien, Frederic Michon

## Abstract

The tear film plays an essential role in corneal protection and regeneration following injury. Although the cornea is a structurally conserved organ across terrestrial vertebrates, the extent to which tear film mediated wound healing responses are evolutionarily conserved remains unclear. This study aimed to identify core and species-specific molecular pathways activated in the tear film during corneal wound healing in humans and mice. We conducted a meta analysis of tear proteomic datasets from human subjects undergoing photorefractive keratectomy (PRK) and mice subjected to mechanical corneal abrasion. Differentially expressed proteins were identified and subjected to Reactome and Gene Ontology (GO) enrichment analyses to determine conserved and divergent biological responses. Approximately one third of the tear film proteomic response to corneal injury was conserved across species. Shared upregulated pathways included complement activation, actin cytoskeletal remodeling, protein synthesis, and acute inflammatory responses. Simultaneously, pathways related to adaptive immunity, proteolysis, and general metabolism were consistently downregulated. Human specific responses were enriched in secretory pathways, vesicle trafficking, and immune surveillance, whereas murine specific responses highlighted mitochondrial activation, oxidative metabolism, and stress adaptation. These distinctions reflect species dependent physiological strategies in managing epithelial repair. Our findings reveal a conserved molecular framework that governs corneal wound healing across species, with notable species specific adaptations. This cross species comparison underscores the translational relevance of tear film analysis and supports the development of targeted therapies tailored to human specific wound healing mechanisms in ocular surface disease.

## Introduction

The vertebrate cornea represents a highly conserved anatomical and physiological structure across diverse species, characterized by transparency, avascularity, and a meticulously organized extracellular matrix crucial for optimal visual performance^1,2^. Despite substantial evolutionary divergence among vertebrates, the fundamental architecture of the cornea—comprising the epithelium, stroma, and endothelium—remains remarkably consistent^3^. This structural conservation highlights strong evolutionary pressures aimed at preserving corneal transparency and integrity, both essential for vision-dependent behaviors such as predation, navigation, and reproduction.

The evolutionary transition from aquatic to terrestrial life necessitated significant physiological adaptations, among which the development of the tear film stands prominent. This multilayered fluid interface is crucial for maintaining corneal hydration, providing trophic support, offering protection against pathogens, and preserving optical clarity^4^. The emergence of the tear film is tightly coupled to terrestrial adaptation, acting as a defense mechanism against new environmental stressors such as desiccation, mechanical damage, and microbial threats, all of which intensified upon terrestrial colonization. Unlike aquatic ancestors whose corneal hydration was inherently supported by their aquatic surroundings, terrestrial vertebrates evolved this sophisticated tear film to actively maintain ocular surface homeostasis.

Remarkably, across terrestrial vertebrate species, the tear film exhibits strikingly conserved structural and functional characteristics. It consists of three distinct yet integrated layers: a superficial lipid layer, an intermediate aqueous layer, and an inner mucinous layer^5^. The lipid layer, predominantly produced by the meibomian glands, prevents evaporation and stabilizes the ocular surface. The aqueous layer, secreted by lacrimal glands, delivers essential nutrients, growth factors, oxygen, and antimicrobial molecules. The mucinous layer, primarily produced by conjunctival goblet cells, ensures stable adhesion of the tear film to the hydrophobic corneal epithelium. The persistent conservation of this trilaminar structure emphasizes its evolutionary robustness as an effective physiological response to terrestrial environmental pressures.

The integrity and functionality of the tear film are closely intertwined with overall corneal health. Disruptions in tear film composition or stability underpin various ocular surface disorders, including dry eye syndrome, corneal ulcers, and neurotrophic keratitis^4,6,7^. Therefore, uncovering novel therapeutic targets within the tear film could profoundly impact clinical strategies, enhancing corneal healing processes and preserving vision. Achieving these therapeutic advances requires an in-depth understanding of tear film dynamics, particularly the molecular alterations occurring in response to corneal injury.

To address this hypothesis, we performed a comparative analysis of tear film proteomic changes during corneal wound healing in murine and human models. We leveraged two pivotal datasets generated in our recent studies: one describing murine corneal healing after physical abrasion^8^, and the other detailing human corneal recovery following photorefractive keratectomy (PRK)^9^. Both proteomic datasets are publicly available, and were obtained on the same mass-spectrometry machinerie, facilitating a direct comparative approach. By re-analyzing these data, we identified conserved molecular mediators that constitute the fundamental response of the tear film to corneal injury. Elucidating this shared molecular framework holds promise for identifying universal therapeutic targets that enhance corneal repair across species and accelerate translational developments for ocular surface diseases.

By identifying these fundamental molecular signatures, our study aims to pinpoint core therapeutic targets with broad applicability across species. Such comparative investigations are expected to expedite the development of innovative therapeutic interventions, offering significant translational potential and paving the way for future research in ocular surface disease management.

## Results

### Description of the tear film composition modulation after corneal wounding

Tear film samples were collected immediately and three days following photorefractive keratectomy (PRK) in humans^9^, and at 6, 12, 18, and 24 hours after mechanical corneal abrasion in mice^8^. We considered all proteomic modulations occurring within these timeframes as indicative of the corneal injury response. The detailed comparative analysis of human and murine proteomic responses is summarized in **Table 1**, and the list of proteins is in **Table S1**. In total, 2025 proteins were identified in human tear samples, while murine samples revealed 3618 proteins. Notably, a similar number of proteins were found to be upregulated in both species (human: 667; mouse: 589). In contrast, the number of downregulated proteins in murine tears (640 proteins) was approximately three-fold higher than in human tears (242 proteins). Consequently, due to the overall lower number of proteins detected in human samples, the proportion of proteins modulated in response to injury was higher in humans. Interestingly, the human tear film response predominantly involved protein upregulation (representing 73.38% of the total injury response), whereas the murine response showed a balanced pattern, with approximately equal proportions of upregulated and downregulated proteins. Finally, a comparative analysis of the protein identities revealed that around one-third of the injury-associated proteomic response was conserved between mice and humans. This substantial overlap underscores a highly conserved core response, alongside significant species-specific proteomic adaptations.

**Table 1.**
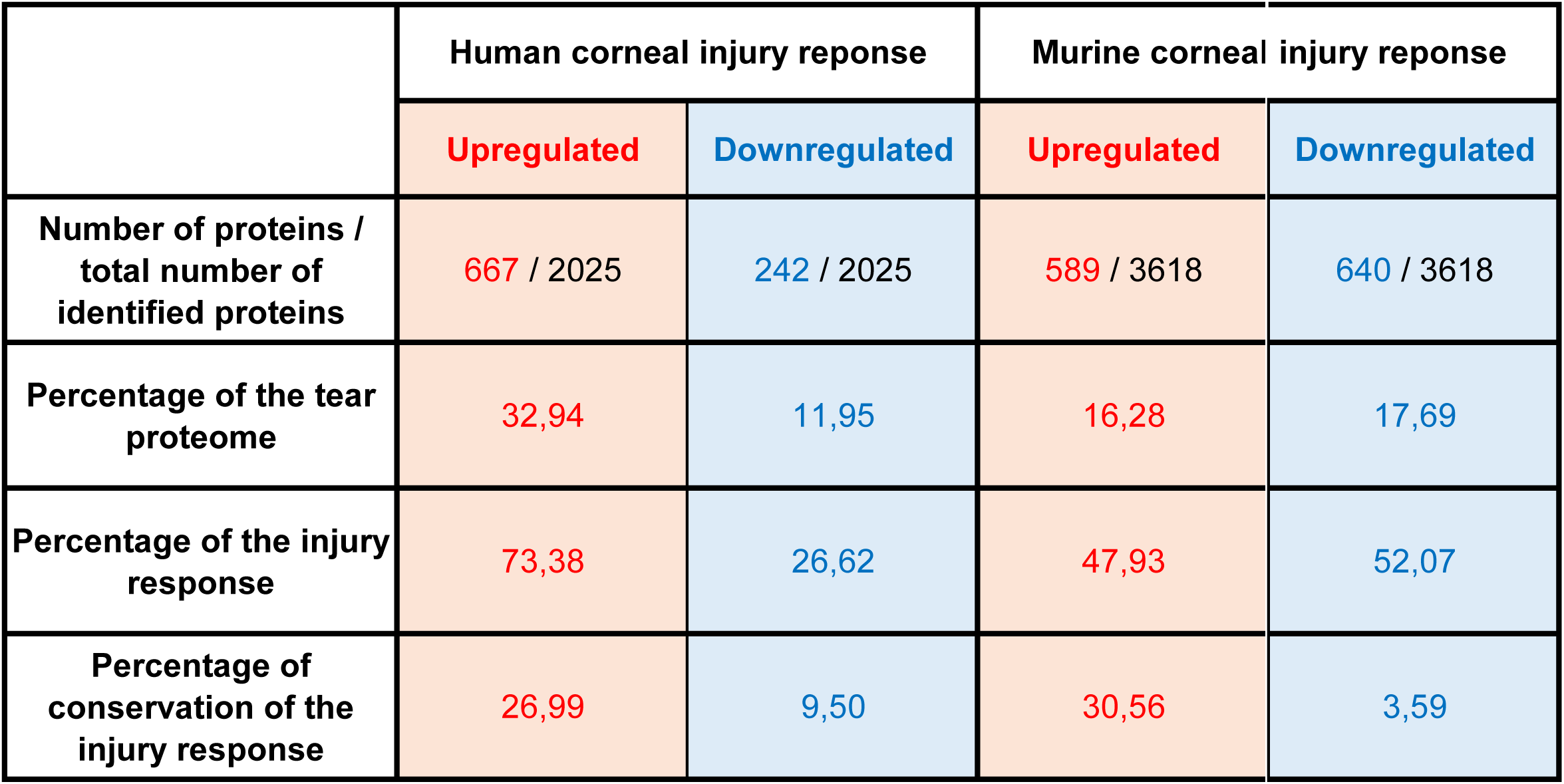
Overview of the corneal injury response in human and mouse.

### Conservation of the identified proteins

We identified 180 proteins that were upregulated and 23 proteins that were downregulated during the corneal injury response in both human and murine models (**Table S2**). To elucidate the functional significance of these differentially expressed proteins, we conducted comprehensive enrichment analyses using Reactome, GO Biological Process, and GO Molecular Function databases, aiming to reveal crucial biological pathways significantly modulated during corneal wound healing (**Table 2**, **Tables S3-S8**).

**Table 2.**
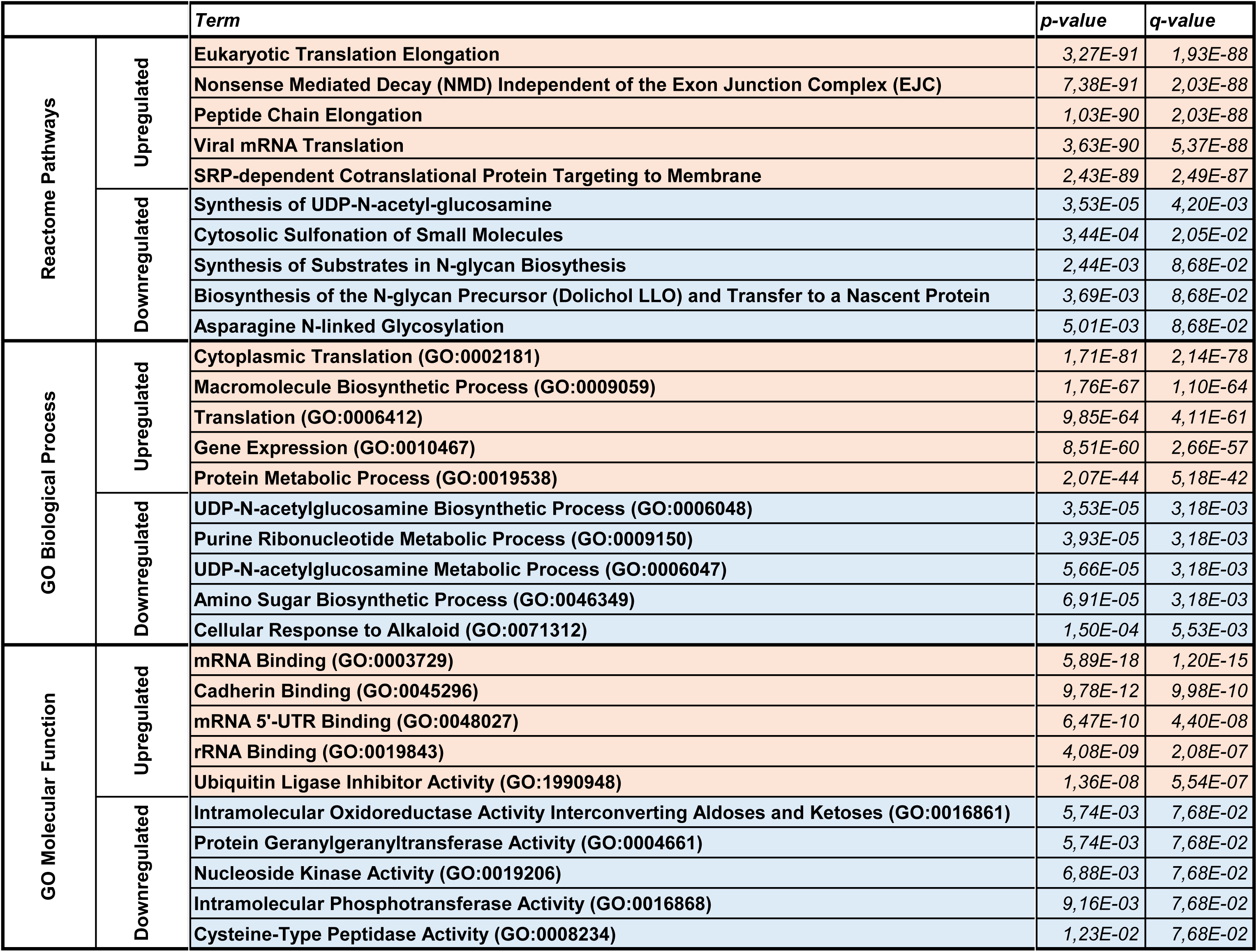
Top 5 *Reactome Pathways*, *GO Biological Process, GO Molecular Functions* terms regulated by the proteins regulated similarly in human and mouse during corneal wound healing.

Our analysis highlighted a predominant upregulation of molecular pathways associated with protein synthesis and translation, specifically eukaryotic translation elongation, nonsense-mediated decay, and viral mRNA translation. Furthermore, the most significantly modulated GO biological processes and molecular functions pointed towards extensive cellular remodeling, adhesion dynamics, and protein regulation, indicating active roles in tissue restructuring and repair during corneal healing (**Table 2**).

Additionally, pathways recognized for their roles in corneal repair, such as axon guidance, keratinization, and Notch signaling activity, were notably upregulated (**Table S3**). Processes critical to effective wound closure, including cell motility (**Table S4**) and fibroblast growth factor (FGF) binding (**Table S5**), also demonstrated significant elevation. Conversely, the limited number of downregulated factors led to suppression of pathways related to axonal growth inhibition (**Table S6**), epithelial cell-cell adhesion (**Table S7**), and actin binding (**Table S8**)—processes essential for the dynamic rearrangement of epithelial cells during wound healing.

While this targeted analysis effectively identified essential biological processes in corneal wound healing, we recognized its inherent limitations due to the complexity and coordination of biological pathways. Therefore, we expanded our investigation by directly analyzing the reactome pathways, GO biological processes, GO cellular components and GO molecular functions modulated in the original human and mouse proteomic datasets to achieve a more holistic understanding of the corneal wound healing response.

### Conservation of integrative functions during corneal wound healing

We extracted the proteins differentially regulated during corneal wound healing in the human (909 proteins identified) and murine (1229 proteins identified) tear film. Then, we analyzed the various integrative functions associated with these factors.

To identify conserved biological processes driving corneal wound healing, we performed pathway enrichment analyses using Reactome and Gene Ontology (GO) databases, focusing on the tear proteome profiles in both human and murine models. We specifically extracted shared upregulated and downregulated terms during the healing response to determine the core molecular programs mobilized during epithelial injury. The results reveal a highly coordinated and evolutionarily preserved response involving innate immune activation, cytoskeletal remodeling, controlled proteolysis, and metabolic reprioritization.

Among the Reactome pathways (**Table 3**, **Table S9**), complement activation emerged as a dominant feature, with strong upregulation of “Activation of C3 and C5” and “Alternative Complement Activation.” These cascades are central to the innate immune defense, mediating pathogen clearance, opsonization, and the recruitment of immune effectors to the injury site. The tear film appears to function as a first-line immune barrier, rapidly responding to epithelial breaches. Notably, this is accompanied by upregulation of the “Acute Inflammatory Response” and “Acute-Phase Response” GO terms, further supporting the role of the tear proteome in delivering immune mediators such as complement proteins, cytokines, and antimicrobial peptides directly to the ocular surface.

**Table 3.**
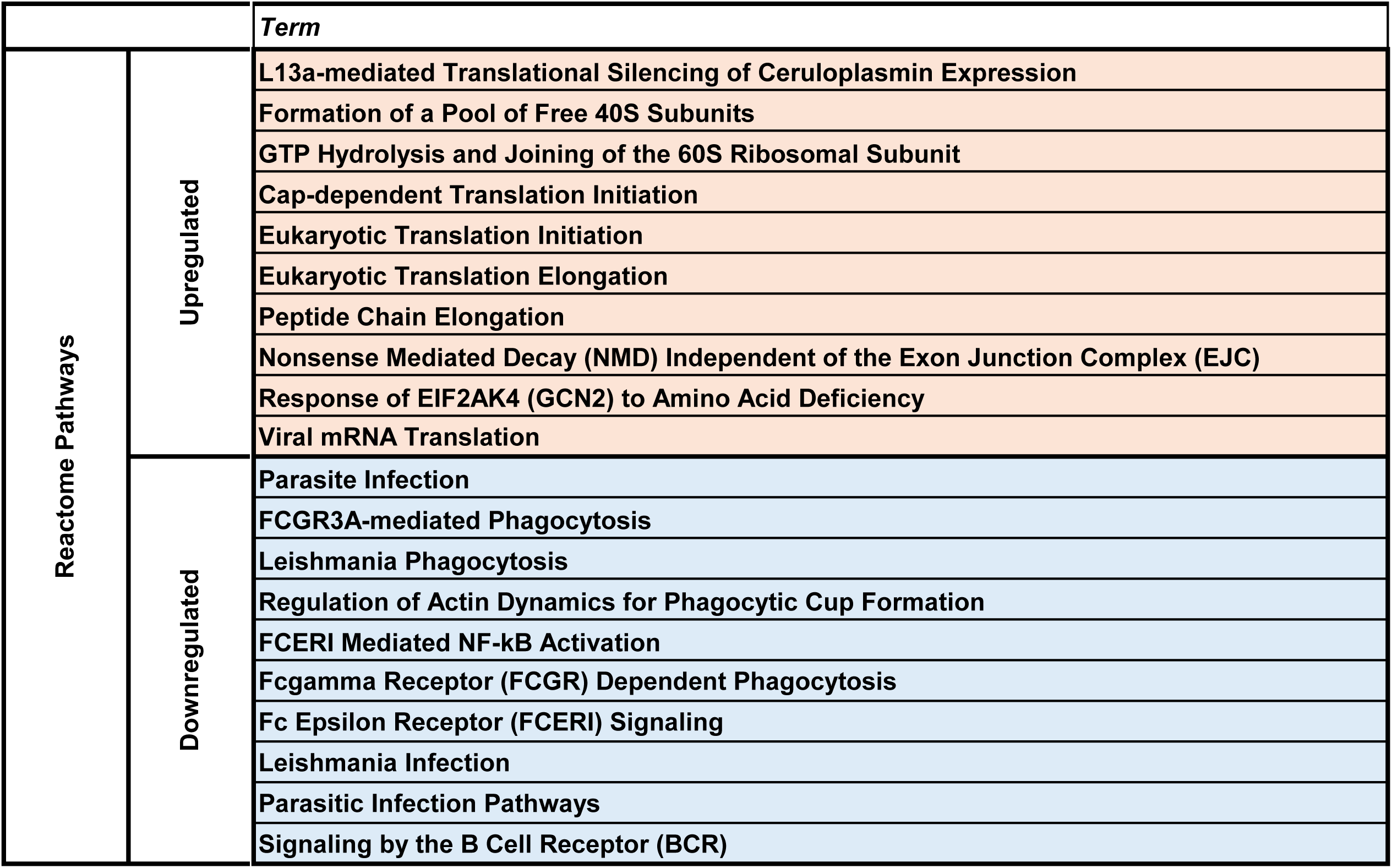
Top 10 *Reactome Pathways* terms regulated similarly in human and mouse during corneal wound healing.

Interestingly, while innate immune pathways were activated, several components of the adaptive immune system, including “MHC Class II Antigen Presentation” and “Fc Receptor Signaling,” were consistently downregulated. This suggests an intentional suppression of adaptive immune mechanisms during the acute wound healing phase, likely to prevent overactivation and subsequent tissue damage. Similarly, Reactome terms related to immunoglobulin glycosylation and antigen processing were also reduced. Such selective immune modulation highlights the unique requirement of the cornea to balance immune defense with the preservation of optical clarity.

In parallel with immune activation, we observed a robust upregulation of pathways associated with cytoskeletal dynamics. GO terms such as “Actin Filament Organization,” “Actomyosin Structure Organization,” and “Adherens Junction” were prominently enriched (**Table 4**, **Table S10**). These processes are central to epithelial cell migration, a critical step in resurfacing the denuded corneal surface. The upregulation of actin and cadherin binding molecular functions further supports a program of collective epithelial cell movement, coordinated by adherens junction remodeling and actomyosin contractility.

**Table 4.** Top 10 *GO Biological Process* terms regulated similarly in human and mouse during corneal wound healing.

|  |  | Term |
| --- | --- | --- |
| GO Biological Process | Upregulated | Cytoplasmic Translation (GO:0002181) |
|  |  | Macromolecule Biosynthetic Process (GO:0009059) |
|  |  | Translation (GO:0006412) |
|  |  | Gene Expression (GO:0010467) |
|  |  | Protein Metabolic Process (GO:0019538) |
|  |  | RNA Processing (GO:0006396) |
|  |  | protein-RNA Complex Assembly (GO:0022618) |
|  |  | mRNA Splicing, via Spliceosome (GO:0000398) |
|  |  | Ribonucleoprotein Complex Biogenesis (GO:0022613) |
|  |  | Ribosomal Small Subunit Biogenesis (GO:0042274) |
|  | Downregulated | Purine Ribonucleotide Catabolic Process (GO:0009154) |
|  |  | Purine Ribonucleotide Metabolic Process (GO:0009150) |
|  |  | Protein Geranylgeranylation (GO:0018344) |
|  |  | Macromolecule Modification (GO:0043412) |
|  |  | UDP-N-acetylglucosamine Biosynthetic Process (GO:0006048) |
|  |  | Protein Prenylation (GO:0018342) |
|  |  | Proteolysis (GO:0006508) |
|  |  | Negative Regulation of Interferon-Beta Production (GO:0032688) |
|  |  | UDP-N-acetylglucosamine Metabolic Process (GO:0006047) |
|  |  | AMP Metabolic Process (GO:0046033) |

On a structural level, cellular components (**Table 5**, **Table S11**) involved in cytoskeletal support and intercellular junctions—such as the actin cytoskeleton and adherens junctions—were enriched, reinforcing the concept that epithelial sheet migration during wound healing relies on the dynamic reassembly of intracellular scaffolds and adhesion complexes.

**Table 5.** Top 10 *GO Cellular Components* terms regulated similarly in human and mouse during corneal wound healing.

|  |  | *Term* |
| --- | --- | --- |
| GO Cellular Components | Upregulated | Focal Adhesion (GO:0005925) |
|  |  | Cell-Substrate Junction (GO:0030055) |
|  |  | Cytosolic Large Ribosomal Subunit (GO:0022625) |
|  |  | Large Ribosomal Subunit (GO:0015934) |
|  |  | Cytosolic Small Ribosomal Subunit (GO:0022627) |
|  |  | Small Ribosomal Subunit (GO:0015935) |
|  |  | Secretory Granule Lumen (GO:0034774) |
|  |  | Intracellular Organelle Lumen (GO:0070013) |
|  |  | Collagen-Containing Extracellular Matrix (GO:0062023) |
|  |  | Ribosome (GO:0005840) |
|  | Downregulated | Vesicle (GO:0031982) |
|  |  | Golgi Membrane (GO:0000139) |
|  |  | Ficolin-1-Rich Granule Membrane (GO:0101003) |
|  |  | Tertiary Granule (GO:0070820) |
|  |  | Secretory Granule Membrane (GO:0030667) |
|  |  | Vesicle Membrane (GO:0012506) |
|  |  | Vacuolar Lumen (GO:0005775) |
|  |  | Ficolin-1-Rich Granule (GO:0101002) |
|  |  | Specific Granule Lumen (GO:0035580) |
|  |  | Bounding Membrane of Organelle (GO:0098588) |

Protease activity, while essential for matrix remodeling and immune defense, must be tightly regulated in the cornea to avoid stromal degradation and opacity (**Table 6**, **Table S12**). In line with this, multiple peptidase functions were significantly downregulated, including “Aminopeptidase Activity,” “Cysteine-Type Peptidase Activity,” and “Calcium-Dependent Phospholipid Binding.” This suggests a conserved strategy to restrain excessive proteolysis during healing, likely protecting the extracellular matrix (ECM) from premature breakdown and preserving corneal transparency.

**Table 6.** Top 10 *GO Molecular function* terms regulated similarly in human and mouse during corneal wound healing.

|  |  | *Term* |
| --- | --- | --- |
| GO Molecular Function | Upregulated | Cadherin Binding (GO:0045296) |
|  |  | mRNA Binding (GO:0003729) |
|  |  | Endopeptidase Inhibitor Activity (GO:0004866) |
|  |  | Serine-Type Endopeptidase Inhibitor Activity (GO:0004867) |
|  |  | Calcium Ion Binding (GO:0005509) |
|  |  | Peptidase Inhibitor Activity (GO:0030414) |
|  |  | Metal Ion Binding (GO:0046872) |
|  |  | Lipoprotein Particle Receptor Binding (GO:0070325) |
|  |  | Actin Binding (GO:0003779) |
|  |  | mRNA 3'-UTR Binding (GO:0003730) |
|  | Downregulated | Calcium-Dependent Phospholipid Binding (GO:0005544) |
|  |  | Transition Metal Ion Binding (GO:0046914) |
|  |  | Exopeptidase Activity (GO:0008238) |
|  |  | Lysophospholipase Activity (GO:0004622) |
|  |  | Cysteine-Type Peptidase Activity (GO:0008234) |
|  |  | Endopeptidase Activity (GO:0004175) |
|  |  | ADP Binding (GO:0043531) |
|  |  | Aminopeptidase Activity (GO:0004177) |

Similarly, terms associated with vesicle-mediated secretion, such as “Golgi Membrane” and “Vesicle,” were downregulated, pointing to a temporary reprioritization away from constitutive secretion. While certain vesicle-related pathways (e.g., COPI trafficking) may be locally upregulated to supply healing factors, broader suppression of secretory machinery may reflect a resource-saving mechanism or a means to limit inflammation.

Despite the downregulation of glycosylation and certain biosynthetic pathways, the tear proteome revealed signs of upregulated protein synthesis, including the activation of “aminoacyl-tRNA ligase activity” and “5S rRNA binding.” This likely reflects the increased translational demands of epithelial cells undergoing proliferation and repair.

Simultaneously, metabolic shifts were evident. GO biological processes involved in amino sugar biosynthesis, glycerolipid catabolism, and AMP metabolism were downregulated, indicating a reallocation of metabolic resources from general cellular maintenance toward tissue regeneration. These changes align with the concept of a metabolic reboot typical of wound-healing tissues, in which cells temporarily divert energy and substrates to support critical repair pathways^10^.

Altogether, the shared enrichment profile observed in both human and murine tear proteomes reveals a core response to corneal injury that integrates innate immune activation, cytoskeletal remodeling, suppression of excessive proteolysis, and a rewiring of metabolic priorities. However, less than 40% of the corneal injury response were conserved between human and mouse, therefore, we investigated the species-specificities of the tear film response to corneal injury.

### Species-specific molecular pathways shaping corneal wound healing dynamics

To uncover divergent mechanisms between humans and mice during corneal wound healing, we analyzed species-specific enrichment profiles derived from the tear proteome. Using Reactome and Gene Ontology annotations, we identified biological processes, cellular components, and molecular functions that were selectively enriched in either human or murine models. These findings reflect distinct physiological priorities and tissue responses, shedding light on both shared and unique aspects of the corneal healing trajectory in different organisms.

The human-specific Reactome enrichment terms (**Table 7**, **Table S13**) revealed a strong emphasis on complement activation and secretory trafficking. Key pathways included the “Complement Cascade,” “Intrinsic Pathway of Fibrin Clot Formation,” and “COPI-mediated Anterograde Transport,” alongside “Golgi-Associated Vesicle Biogenesis” and “RAB Geranylgeranylation.” This enrichment pattern indicates a coordinated mobilization of secretory and immune machinery in the human tear film. The tear proteome appears to serve not only as a reservoir of inflammatory mediators but also as a conduit for vesicle-mediated replenishment of protective and structural proteins essential for healing. Concordantly, human-specific GO Biological Processes were enriched in “Protein Localization,” “Positive Regulation of Cell-Substrate Adhesion,” and “Positive Regulation of Secretion by Cell,” emphasizing directed trafficking and epithelial migration (**Table 8**, **Table S14**). The upregulation of endothelial barrier formation and ERK/MAPK signaling cascades further supports the notion of a complex, tightly regulated epithelial repair program involving both secretory and migratory components. At the cellular level, human-enriched terms included “Cytoplasmic Vesicle Membrane,” “Very-Low-Density Lipoprotein Particles,” and “Membrane Attack Complex,” reflecting not only vesicular dynamics but also immune surveillance functions (**Table 9**, **Table S15**). The presence of lipoprotein particle-related terms also suggests modulation of lipid-based components of the tear film, which are essential for ocular surface stability. Human-specific molecular functions highlighted an increase in “GTPase Activity,” “GDP Binding,” and “Myosin Binding,” which point to cytoskeletal control and vesicle trafficking via small GTPase signaling (**Table 10**, **Table S16**). The actin-myosin axis appears especially prominent, likely supporting epithelial cell shape changes and dynamic migration.

**Table 7.**
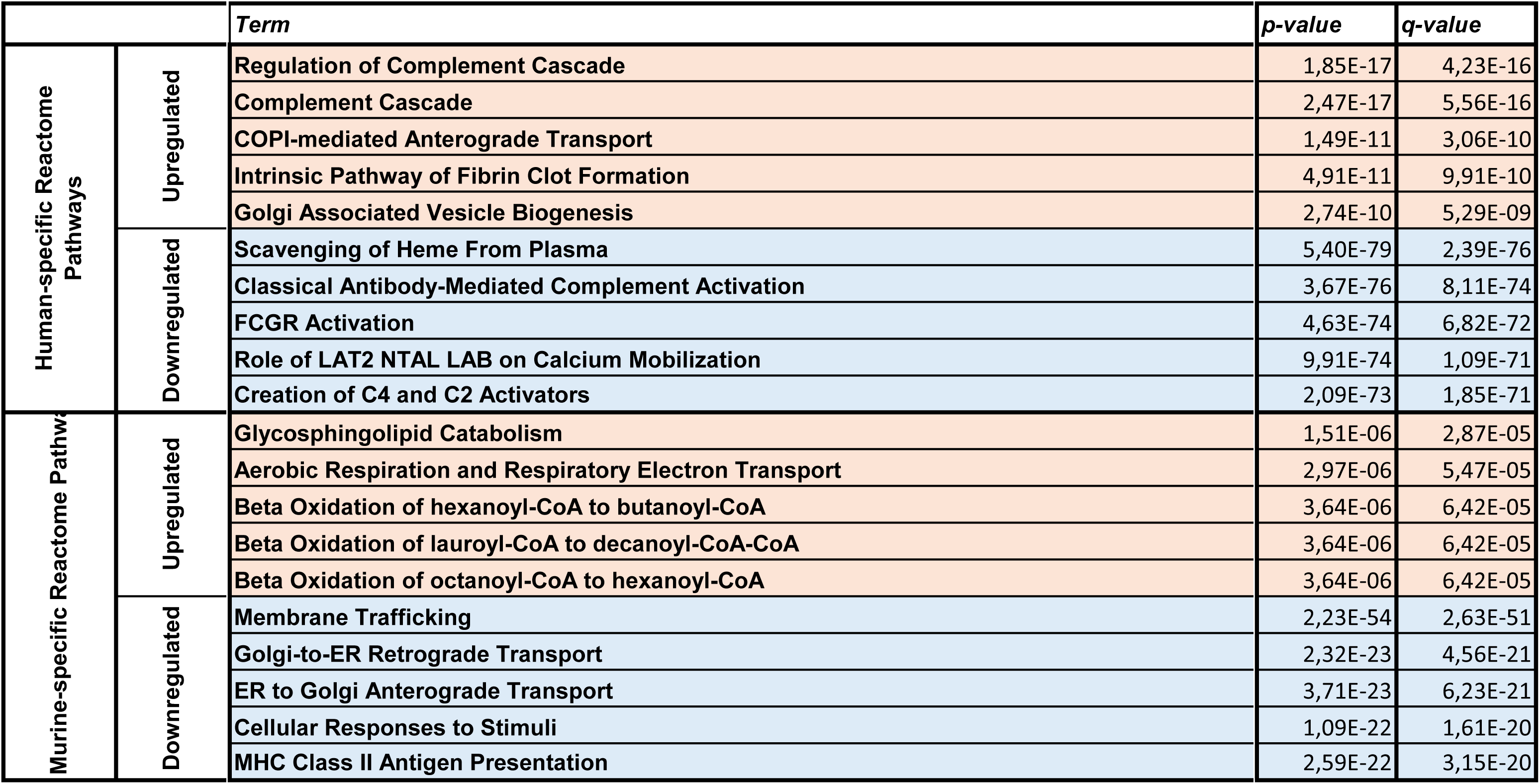
Top 5 species-specificities in *Reactome Pathways* regulation during corneal wound healing.

**Table 8.**
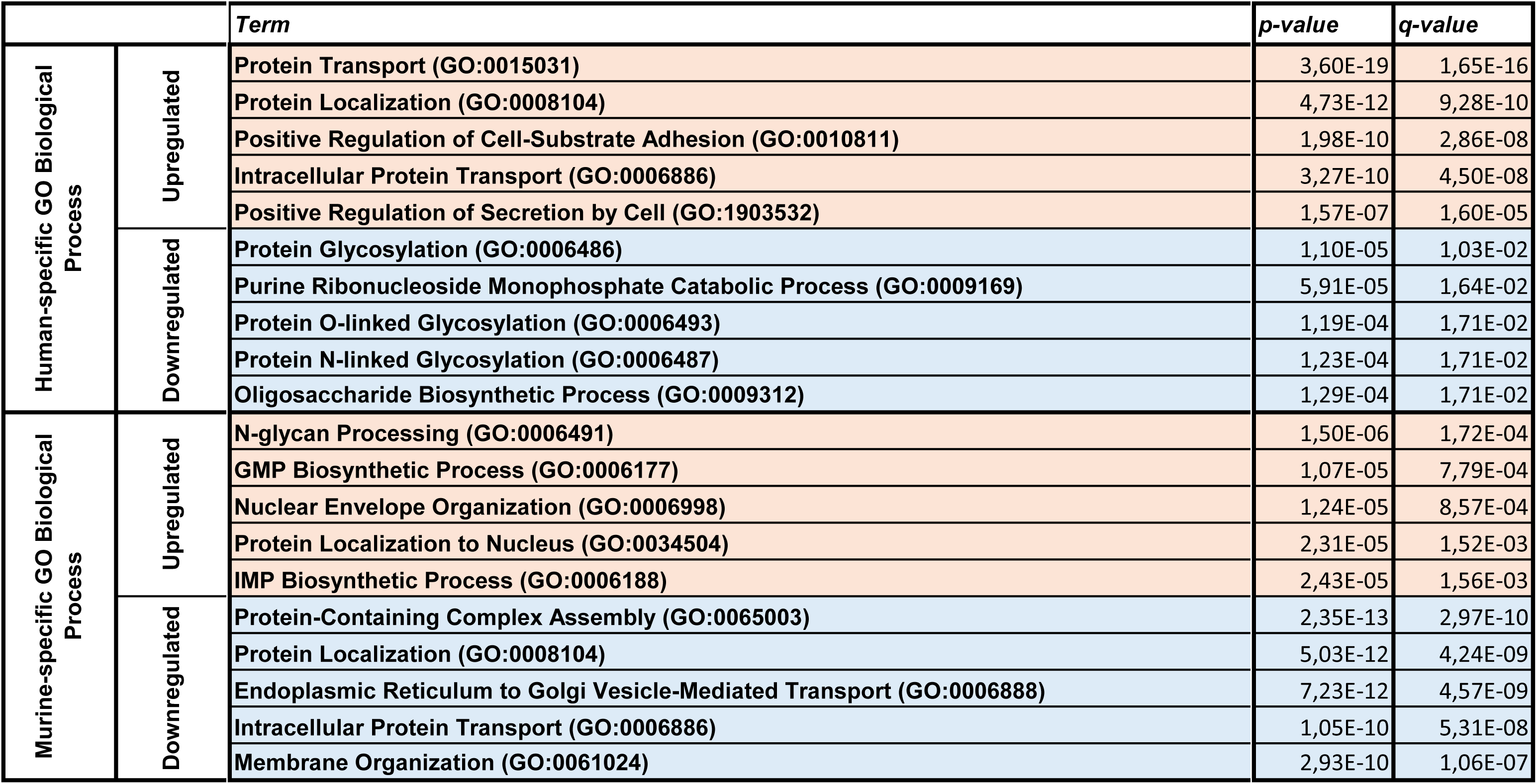
Top 5 species-specificities in *GO Biological Process* regulation during corneal wound healing.

**Table 9.**
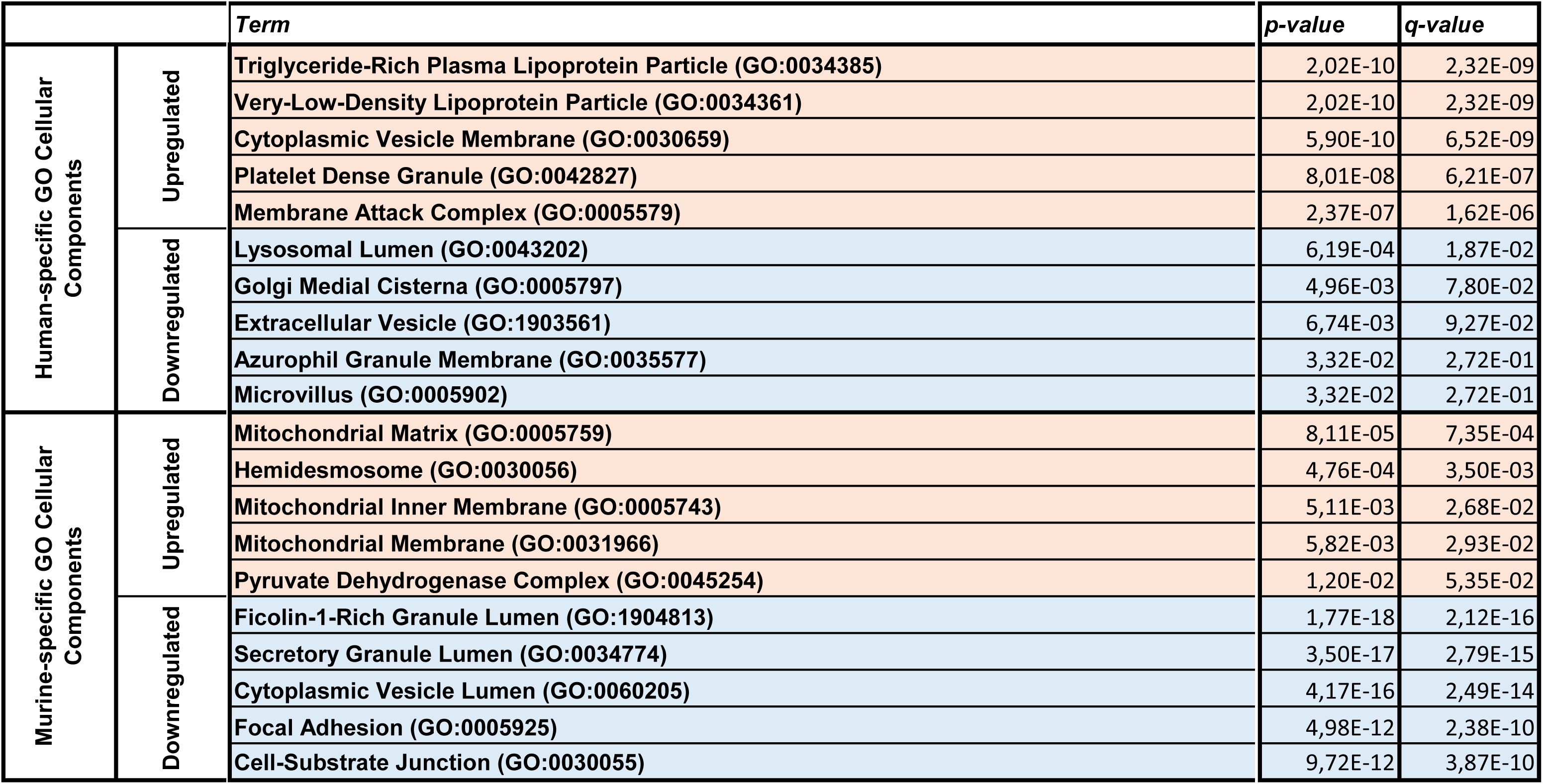
Top 5 species-specificities in *GO Cellular Components* regulation during corneal wound healing.

**Table 10.**
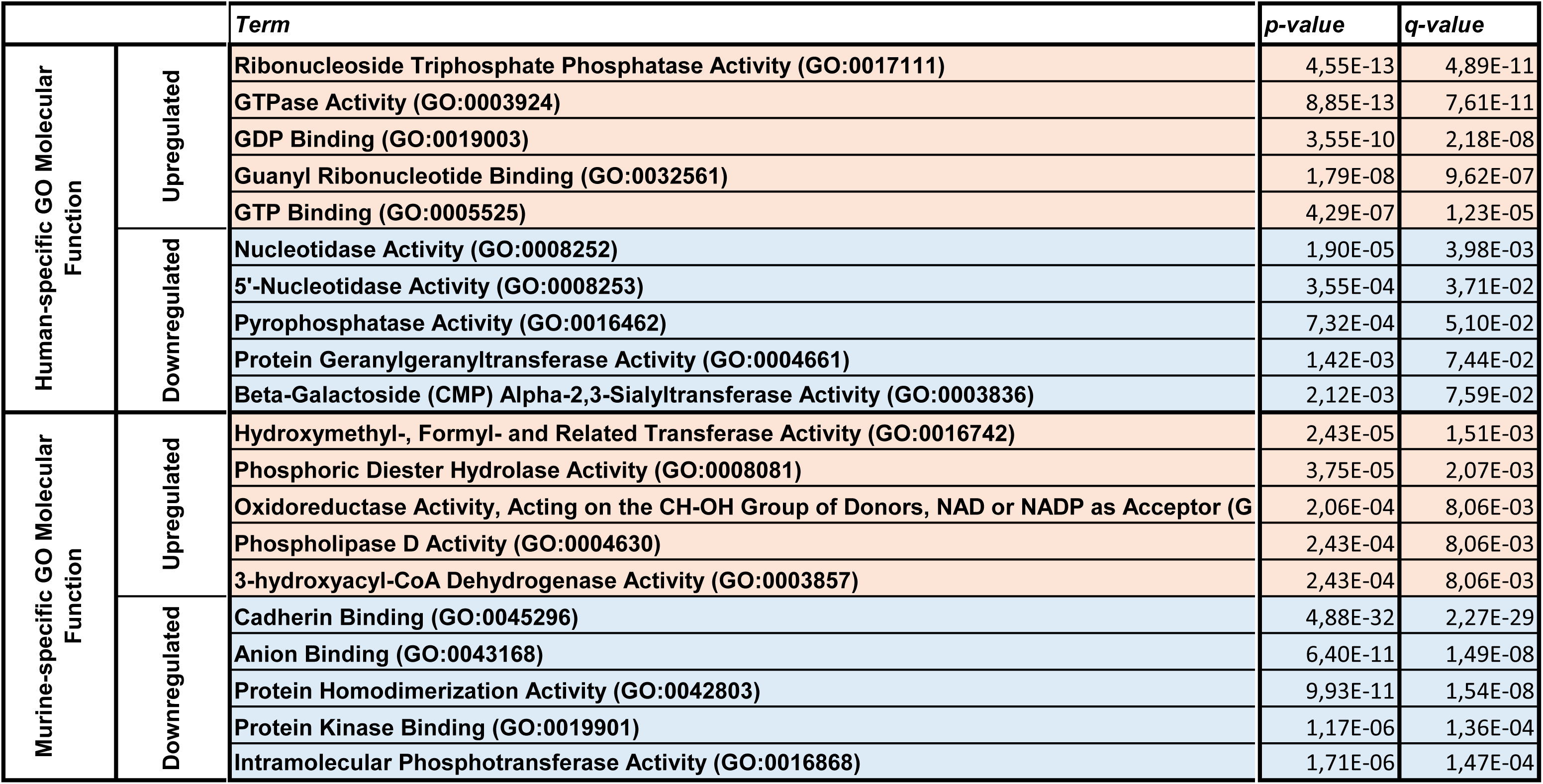
Top 5 species-specificities in *GO Molecular Function* regulation during corneal wound healing.

Together, these results suggest that the human cornea relies on a secretory-driven and immunologically alert environment, with a heavy reliance on vesicle-mediated trafficking and barrier integrity maintenance. This response likely reflects evolutionary pressure to protect a larger, longer-lived, and exposed corneal surface from both microbial invasion and physical desiccation.

In contrast, murine-specific enrichment profiles were dominated by mitochondrial and metabolic processes. Reactome terms such as “Aerobic Respiration and Respiratory Electron Transport,” and various “Beta Oxidation” pathways revealed a pronounced shift toward oxidative energy production and lipid metabolism (**Table 7**, **Table S13**). These pathways likely support the intense energy demands of rapid re-epithelialization, a hallmark of the murine corneal response. GO Biological Processes enriched in the murine tear proteome included “GMP Biosynthetic Process,” “Nuclear Envelope Organization,” and “Unfolded Protein Response,” consistent with a metabolically activated and stress-adapted cellular state (**Table 8**, **Table S14**). Upregulation of nucleotide biosynthesis suggests preparation for proliferation and DNA replication, while nuclear reorganization and UPR activation indicate mechanisms for coping with translational load and oxidative stress. At the structural level, murine-enriched GO Cellular Components centered on mitochondria: “Mitochondrial Inner Membrane,” “Pyruvate Dehydrogenase Complex,” and “Alpha-Ketoacid Dehydrogenase Complex” all point toward active mitochondrial energy metabolism (**Table 9**, **Table S15**). In parallel, enrichment of “Hemidesmosomes” underscores the reinforcement of epithelial–basement membrane anchoring, stabilizing the newly formed epithelium during high-turnover repair. Molecular functions further reinforce this metabolic theme (**Table 10**, **Table S16**). Enriched murine-specific activities included “Oxidoreductase Activity,” “Phospholipase D Activity,” and “3-Hydroxyacyl-CoA Dehydrogenase Activity,” which contribute to lipid breakdown, redox balance, and mitochondrial fatty acid metabolism. These processes are critical for fueling the epithelial regenerative process and maintaining homeostasis under injury-induced oxidative stress.

Collectively, the murine corneal healing response appears to emphasize rapid, energy-intensive, and metabolically efficient tissue regeneration, supported by mitochondrial activation and reduced reliance on external secretory inputs. This contrasts with the human reliance on tear-based immune and secretory support and reflects species-specific strategies shaped by corneal size, life span, and immune architecture.

## Discussion

The present meta-analysis of tear proteomic dynamics during corneal wound healing reveals an evolutionarily conserved yet distinctly modulated biological response across species. By comparing murine and human datasets^8,9^, we identified not only shared molecular mediators underpinning the core wound-healing program, but also species-specific adaptations that reflect divergent physiological priorities. These results offer both mechanistic insights into corneal regeneration and a rational basis for developing targeted therapeutic strategies adapted to human biology.

Our initial observation was a notable difference in the number of proteins identified: 2,025 in human tear film versus 3,618 in murine samples. This discrepancy likely stems from differences in sampling methodologies. Human tears were collected using Schirmer strips, whereas murine tears were obtained via glass microcapillary. Previous studies have demonstrated that while capillary collection offers superior reproducibility compared to Schirmer strips, it generally yields a lower protein concentration and reduced diversity^11,12^. Consequently, the higher number of proteins detected in murine samples suggests that the murine tear film may possess intrinsically greater proteomic complexity.

At the core of the conserved response is a tightly regulated interplay between innate immunity, cytoskeletal remodeling, proteolysis suppression, and metabolic reprioritization. The early upregulation of complement activation pathways—particularly the alternative pathway (C3/C5) and terminal lytic components—emphasizes the tear film’s role as a frontline immune barrier. This innate immune activation likely provides rapid antimicrobial defense, neutralizing pathogens introduced through corneal abrasion or surgery. Simultaneously, the suppression of adaptive immune signatures such as MHC class II antigen presentation and Fc receptor signaling suggests a protective strategy to minimize inflammation-driven tissue damage. This dichotomy—immune activation without overactivation—may be especially critical for preserving corneal transparency.

Parallel to immune regulation, our analyses uncovered the robust activation of cytoskeletal pathways, including actin filament organization, actomyosin contractility, and cadherin-mediated adhesion. These pathways are essential for the collective migration of epithelial cells that reseal the wounded surface. Structural components such as adherens junctions and the actin cytoskeleton were strongly enriched, supporting the idea that mechanical integrity and dynamic motility are orchestrated in concert. Notably, this remodeling appears coordinated with increased protein synthesis, as evidenced by the upregulation of tRNA ligase activity and ribosomal binding functions—molecular hallmarks of heightened cellular activity during tissue regeneration.

The regulation of proteolytic activity emerged as another conserved axis. While certain matrix-degrading enzymes may assist in clearing damaged ECM and facilitating migration, our data show consistent downregulation of cysteine-type and aminopeptidase activities. Such restraint is likely vital for protecting stromal collagen from degradation and maintaining the biomechanical integrity of the cornea. In addition, suppression of vesicle-mediated secretion and glycosylation pathways suggests a temporary downshifting of secretory processes, perhaps to reduce metabolic load or avoid excessive extracellular signaling.

Importantly, our findings also highlight species-specific differences that must be carefully considered when translating preclinical data to human contexts. In humans, the tear film response is characterized by pronounced secretory and immune surveillance functions. Complement activation, vesicle trafficking (COPI/Golgi), and endothelial barrier maintenance are central components. These likely reflect both the architectural complexity of the human cornea and its prolonged exposure to environmental insults. The reliance on secretory machinery may represent an evolved adaptation to sustain ocular surface integrity over a longer lifespan and within a more variable ecological niche.

By contrast, the murine cornea responds with a metabolic reboot marked by mitochondrial activation, β-oxidation, and nucleotide biosynthesis. These features support a highly energy-demanding regenerative process, consistent with the rapid epithelial turnover observed in rodents. Furthermore, the enrichment of nuclear envelope reorganization and unfolded protein response pathways underscores the need for cellular resilience during high-throughput protein production. The murine corneal epithelium appears optimized for rapid restoration rather than long-term maintenance, a distinction likely shaped by differences in size, ocular exposure, and immune architecture.

These divergent signatures underscore the limitations of directly extrapolating murine data to human therapeutics. While the murine model remains invaluable for delineating fundamental biological processes, its metabolic-heavy profile suggests it may disproportionately favor strategies that enhance energy metabolism or proliferation. In contrast, human-centered therapies may benefit from modulating secretory pathways, immune resolution mechanisms, or cytoskeletal stabilization. Our comparative approach provides a rational framework for tailoring therapeutic design to species-specific biological contexts, thereby improving translational relevance.

In conclusion, this study presents the first integrative, cross-species analysis of tear proteomic signatures during corneal wound healing. It defines a conserved biological core essential for epithelial regeneration and reveals evolutionary adaptations that fine-tune the repair strategy to species-specific constraints. These insights provide a foundation for precision medicine approaches in ophthalmology, enabling the development of therapies that not only accelerate wound closure but also preserve long-term corneal function and visual acuity. Future investigations should explore how these pathways are modulated in pathological states, such as neurotrophic keratitis or diabetic keratopathy, where healing is impaired. Ultimately, a deeper understanding of these molecular programs will illuminate new avenues for restoring and protecting vision across diverse patient populations.

## Material and Methods

### Analysis strategy

We extracted the lists of proteins identified in the murine^8^ and human^9^ tear film during corneal wound healing. We selected the proteins exhibiting a fold change <0.5 or >2, and significantly dysregulated (p-value<0.05). We identified the proteins which were identified in the murine and human tear film.

### Functional and pathway enrichment analyses

We selected the proteins enriched or depleted at each timepoint based on a *p-value*<0.05, and a fold change >2 (enriched), or <0.5 (depleted). The resulting lists of proteins were analyzed using the Enrichr platforme (https://maayanlab.cloud/Enrichr/), followed by Appyter (https://appyters.maayanlab.cloud/) for enrichment analysis visualization. We gathered the analyses related to the Reactome Pathways, GO Biological Pathways, GO Cellular Components, and GO Molecular Function.

## Author contributions

Conceptualization: N.F., M.D., V.D., F.M.; Methodology: N.F., M.D., J.V., C.H., V.D., F.M.; Validation: C.H., F.M.; Formal analysis: N.F., M.D.; Data analysis: N.F., M.D., F.M.; Writing: N.F., M.D., J.V., C.H., V.D., F.M.; Supervision: F.M.; Project administration: F.M.; Funding acquisition: F.M.

## Acknowledgments

This research was supported by ATIP-Avenir program, Inserm, the CHU Montpellier, the Région Occitanie, ANR (ANR-21-CE17-0061, TeFiCoPa), FRM (REP202110014140), Support for research: I-SITE 2024 - program of excellence of the University of Montpellier, CBS2 Doctoral School.

## Supplementary Material

**Table S1. Lists of all proteins up and downregulated in the murine and human tear film during corneal wound healing, in alphabetical order.**

**Table S2. Lists of the proteins that are similarly up and downregulated in the murine and human tear film during corneal wound healing, in alphabetical order.**

**Table S3. List of all the *Reactome Pathways* terms obtained from the proteins that are upregulated similarly in murine and human tear film during corneal wound healing.**

**Table S4. List of all the *GO Biological Process* terms obtained from the proteins that are upregulated similarly in murine and human tear film during corneal wound healing.**

**Table S5. List of all the *GO Molecular Function* terms obtained from the proteins that are upregulated similarly in murine and human tear film during corneal wound healing.**

**Table S6. List of all the *Reactome Pathways* terms obtained from the proteins that are downregulated similarly in murine and human tear film during corneal wound healing.**

**Table S7. List of all the *GO Biological Process* terms obtained from the proteins that are downregulated similarly in murine and human tear film during corneal wound healing.**

**Table S8. List of all the *GO Molecular Function* terms obtained from the proteins that are upregulated similarly in murine and human tear film during corneal wound healing.**

**Table S9. List of all *Reactome Pathways* terms similarly regulated in human and murine tear film, in alphabetical order.**

**Table S10. List of all *GO Biological Process* terms similarly regulated in human and murine tear film, in alphabetical order.**

**Table S11. List of all *GO Cellular Components* terms similarly regulated in human and murine tear film, in alphabetical order.**

**Table S12. List of all *GO Molecular Function* terms similarly regulated in human and murine tear film, in alphabetical order.**

**Table S13. List of all species-specific *Reactome Pathways* terms regulated in human or murine tear film.**

**Table S14. List of all species-specific *GO Biological Process* terms regulated in human or murine tear film.**

**Table S15. List of all species-specific *GO Cellular Components* terms regulated in human or murine tear film.**

**Table S16. List of all species-specific *GO Molecular Function* terms regulated in human or murine tear film.**

